# Lambda Theta Reflectometry: a new technique to measure optical film thickness applied to planar protein arrays

**DOI:** 10.1101/2025.03.26.645463

**Authors:** Alanna M. Klose, Joseph D. Katz, Robert Boni, David Nelson, Benjamin L. Miller

## Abstract

Quantitative protein measurements provide valuable information about biological pathways, immune system functionality, and mechanisms of disease. The most accurate methods for detecting proteins are label-free and preserve native protein binding interactions. Label-free biomolecular interaction analysis includes reflectometry, a group of techniques that detect proteins by measuring the reflectance properties of a thin film on a substrate. Most of these techniques are limited in some way by instrument complexity, sensitivity, or consumable manufacturing requirements. To address these issues, we introduce Lambda Theta Reflectometry (LTR), a new reflectometric technique that measures changes in film thickness by determining the location of null reflectivity as a function of wavelength (lambda) and angle of incidence (theta). The substrate is simultaneously illuminated with a range of angles and wavelengths and reflected light is angularly and spectrally resolved. Our prototype LTR reflectometer can measure SiO_2_ layer thickness with milli-Ångstrom precision. LTR measurements of Si/SiO_2_ oxide films are in excellent agreement with spectroscopic ellipsometry for film thicknesses ranging from 1390-1465 Å. This technique enables sensitive measurements across a range of biological analyte concentrations without requiring stringent control over probe deposition thickness or substrate manufacturing.

---

Biosensors detect and quantify proteins, nucleic acids, and small molecules, and are tools for advancing research and guiding medical treatment.^1^ Label-free biosensors directly measure the interaction of an analyte with a transducer.^2^ The field of label-free optical biosensors^3^ includes surface plasmon resonance^4^, Raman Spectroscopy^5^, integrated photonic devices^6^, and reflectometry^7^. Reflectometry is a group of techniques that measure the optical thickness of a functionalized thin film by illuminating the substrate and evaluating the properties of the reflected light. As light encounters a thin film layer stack, reflections with the layer boundaries interfere with each other encoding information about layer properties onto the reflected light. Biological binding events, such as a protein binding to its receptor, or an antibody binding to its antigen, cause Ångstrom level increases in film thickness. Reflectometric biosensors detect these events by observing the characteristics of the reflected light such as polarization state, spectral interference patterns, or reflectivity at fixed angle or wavelength. Ellipsometry^8^ measures changes in the polarization of reflected light. Oblique-Incidence Reflectivity Difference (OI-RD) microscopy^9^ measures the reflectivity difference between p-polarized and s-polarized light. Reflectometric Interference Spectroscopy (RIfS)^10^ and Biolayer Interferometry (BLI)^11^ illuminate a thin film stack at normal or near normal incidence and evaluate changes in the interference pattern of reflected light as a function of wavelength. Other techniques measure a change in the reflected light intensity at a single or a few discrete wavelengths at fixed angle (Spectral Reflectance Imaging Biosensor (SRIB/IRIS)^12^, 1λ Reflectometry^13^, and Arrayed Imaging Reflectometry (AIR) ^14^). AIR is unique among these techniques in that it leverages the steep drop in reflectivity at an absolute antireflective condition.

An absolute antireflective condition occurs when light waves reflecting off the boundaries of the thin film meet with equal amplitude and opposite phase such that the sum of reflected waves equals zero. An example using a commonly employed material stack consisting of silicon, SiO_2_, and air is shown in Figure 1a. When waves of equal amplitude meet, total destructive interference occurs when the optical path length of light traveling through layer 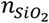 is exactly a half integer multiple of the wavelength of incident light. With the appropriate selection of illumination polarization, wavelength, angle of incidence (AOI), and layer thickness, a condition of zero reflectance can theoretically be achieved (Figure 1b). When optical thickness 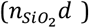 changes, the antireflective condition exists at a different combination of illumination wavelength and AOI.

**Figure 1.**
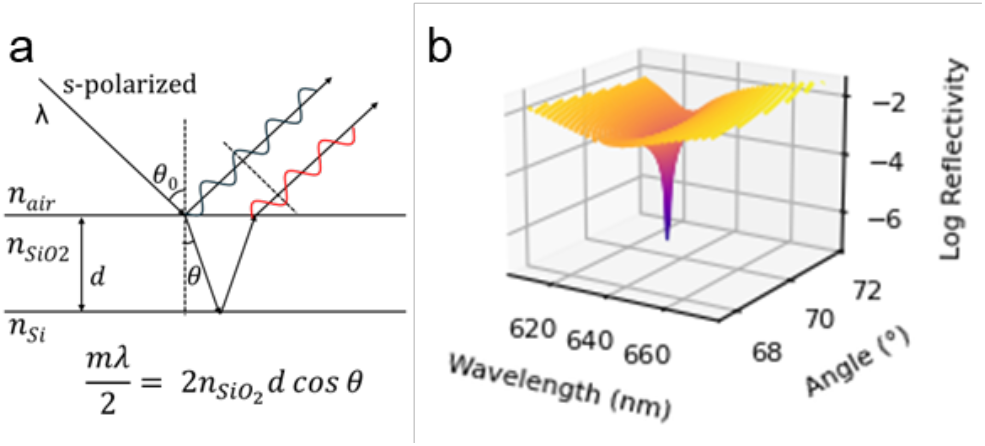
(a) The null reflectivity condition is a function of incident light angle, wavelength, and film thickness. (b) A modeled null condition with a steep drop in reflectivity exists at a specific combination of wavelength and AOI for a thin film with a given optical thickness

The first iteration of AIR, then known as Reflective Interferometry, employed wavelength scanning of s-polarized light at 70.6° angle of incidence to measure reflectivity vs wavelength at several wavelengths around the expected minimum.^15^ A parabolic fit was used to identify the wavelength at minimum reflectivity, from which thickness was calculated using the Fresnel equations. Limitations of this initial design included poor collimation, low s:p polarization ratio, data collection times up to 3 hours per sample, limited spectral resolution and reduced measurement contrast for large changes in layer thickness relative to the initial optimized null condition.

Following generations of AIR simplified the design using a fixed wavelength, 632.8 nm, HeNe laser and included improved imaging optics to measure analyte binding via increased reflectance from an antireflective condition^14^. The relationship between the measured AIR signal and the substrate thickness is established empirically via cross calibration to ellipsometry and is represented by a parabolic fit. As such, layer thickness is not uniquely determined by the data for conditions that deviate from the reflectance null.

The current version of AIR is most accurate only if the thicknesses of the starting SiO_2_ film and adhesion chemistry (typically a silane carrying an electrophilic moiety enabling covalent attachment of probes) are controlled precisely, and probes are deposited at the optimal thickness required to produce an antireflective condition prior to analyte capture.^16^

In the context of biosensing, the thin film is functionalized with probes that have specific affinity to the analyte of interest. Upon binding of analyte, the average thickness of the layer increases according to fractional coverage of binding sites.^17^ In practice, for higher-plex arrays, when many different probe proteins are deposited in different positions on a monolithic substrate, the required control over baseline thickness at each site is difficult to achieve. Errors in probe site starting thickness introduce several possible scenarios that result in reduced measurement accuracy using AIR technique. (Figure S1)

This limitation of AIR motivated the design of a new reflectometric instrument that decouples measurement accuracy from baseline probe thickness while maintaining the steep slope of the antireflective condition enabling highresolution measurements.

Lambda Theta Reflectometry (LTR) addresses the challenges of AIR by illuminating the substrate surface with a range of AOIs and wavelengths to ensure that the null reflectance condition is sampled regardless of initial substrate conditions. By locating the coordinates of the reflectance null in wavelength and angle space, a unique determination of the substrate thickness can be made with milli-Ångstrom precision. The LTR instrument (Figure 2a) measures the reflectivity of a functionalized Si/SiO_2_ substrate with continuous angular and spectral resolution spanning 68°-74° and 590 nm-670 nm respectively. This is accomplished by simultaneously illuminating the substrate with a range of angles and wavelengths using a fiber coupled white light lamp. The output of the lamp is collimated, polarized, and re-imaged on the tilted substrate surface resulting in an elliptical illumination region with major and minor axes of 150 μm and 50 μm. S-polarized light was selected to provide the appropriate reflection amplitudes for fully destructive interference.

**Figure 2.**
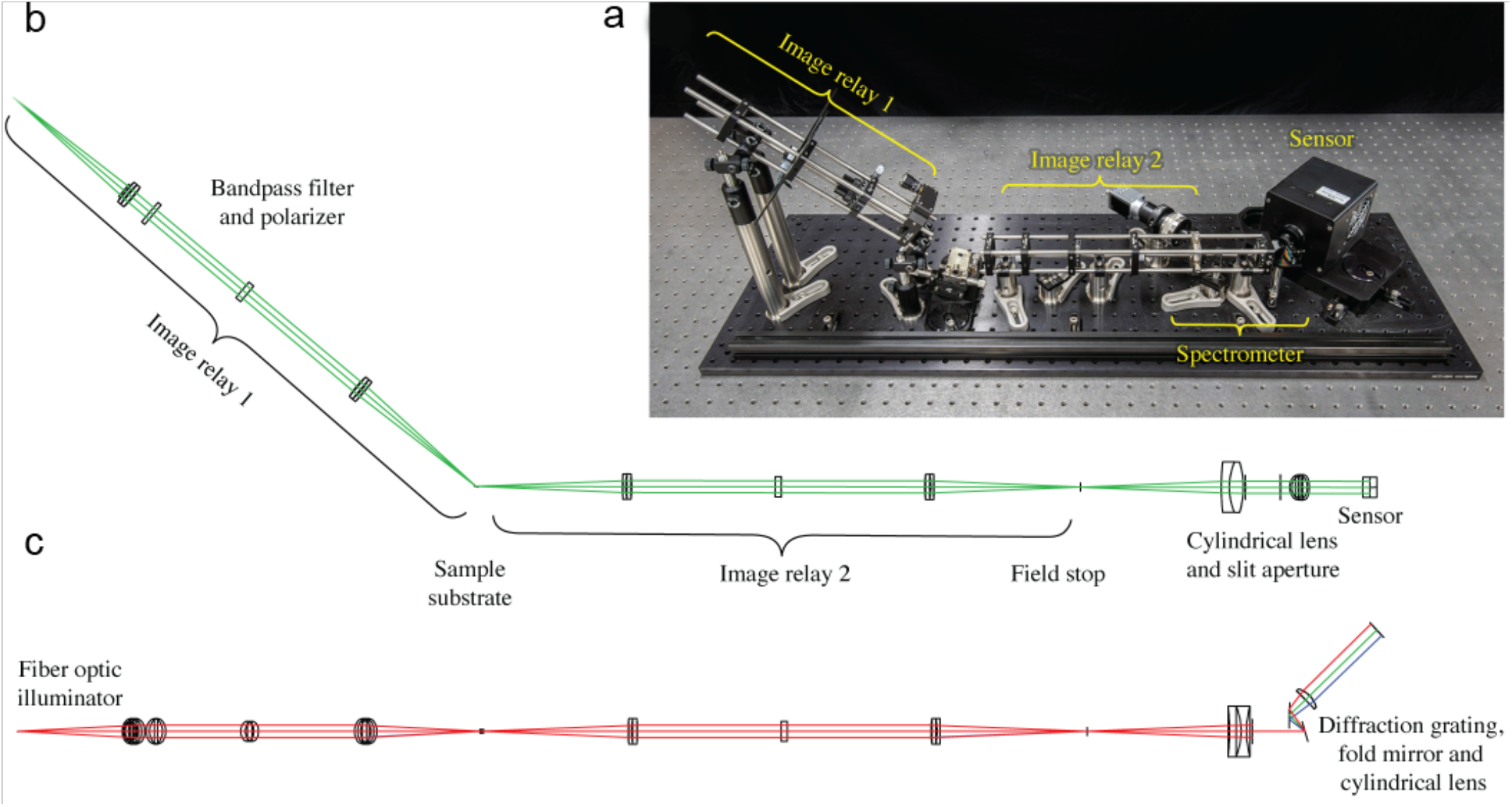
LTR angularly and spectrally resolves the surface reflectance of a sample simultaneously without any moving parts by dispersing the light onto a CCD detector. (a) A computer aided design rendering of the instrument with two orthogonal projections of the optical layout are shown. (b) A side-on view displays the ray fans that are angularly resolved. The range of angles that are sampled (68-74°) is determined by the sample holder tilt angle and the numerical aperture of the illumination image relay. (c) A top-down view displays the ray fans that are spectrally resolved. A bandpass filter limits the wavelengths sampled to (590-670nm). A slit aperture rejects rays that are incident on the sample at compound angles relative to the angular resolution plane. A diffraction grating disperses the remaining rays as a function of wavelength.

Cylindrical optics, a slit aperture, and a diffraction grating are used to orthogonally disperse the reflected light based on the AOI (Figure 2b) and the illumination wavelength (Figure 2c) producing a two-dimensional image that is captured using a CCD camera. The resulting reflectivity data is related to the average thickness of the illumination region. Moving the substrate relative to the fiber core image allows different locations on the substrate surface to be sampled. A secondary alignment camera views the surface to facilitate alignment of the illumination region to individual probe sites deposited on the substrate.

## Results and Discussion

To demonstrate functionality of the LTR technique, we first measured the oxide thickness of a clean Si/SiO_2_ substrate. Each pixel in the experimental LTR image corresponds to a reflectance measurement at a particular wavelength and AOI (Figure 3a). The data reduction process returned film thickness by identifying the conditions of best fit between modeled and measured reflectivity over an entire LTR image. Theoretical reflectance over a range of wavelengths and angles was calculated as a discrete function of film thickness using the characteristic matrix method for multilayered thin films^18^. The theoretical reflectivity of pure s-polarized light incident on an atomically smooth and flat substrate drops to zero, but finite spectral and angular resolution, scattered light at the detector plane, and thin film substrate imperfections limit the reflectivity depth of what can practically be observed. The typical minimum measured reflectivity for our Si/SiO_2_ substrates was ∼5E-4. To account for the deviation in null depth from the theoretical zero, the modeled values were adjusted by adding the minimum reflectivity in the measured data at all points in the modeled data. Three potential sources were considered that could contribute to the experimentally observed null depth. These were s:p polarization ratio, substrate roughness, and scattered light. Reducing the modeled polarization s:p contrast ratio from what was expected when using a Glan-Thompson calcite polarizer from 100,000:1 to 50,000:1 was not sufficient to account for the full difference in minimum reflectivity.

**Figure 3.**
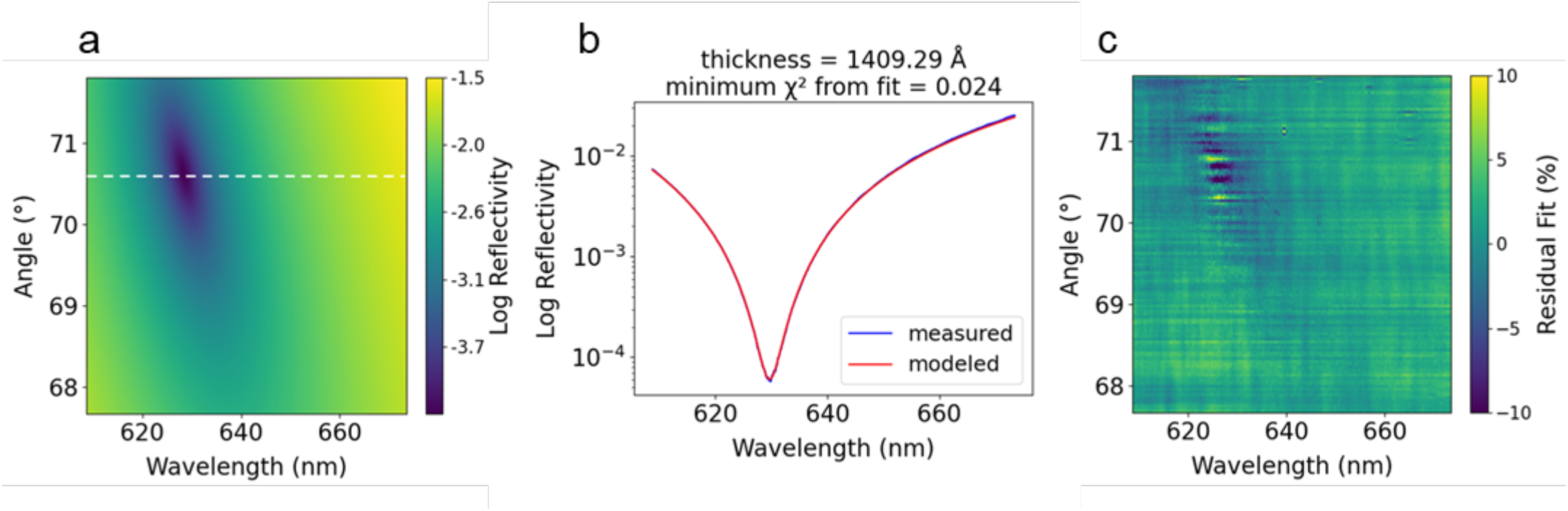
(a) Experimental LTR image with lineout through the null reflectivity condition. (b) Reflectivity vs. wavelength for measured and modeled data at the minimum fit FOM. (c) Residual fit % between measured and modeled reflectivity data at the minimum FOM.

We investigated the substrate surface roughness using atomic force microscopy (AFM) to profile a 4 µm^2^ region of a Si/SiO_2_ substrate (Figure S2). Power spectral density analysis of the AFM micrograph identified low frequency variations in surface height comparable to LTR illumination wavelengths, which could cause a superposition of reflectance properties on the sensor. We simulated this by applying a moving average of thickness values over a 2 nm window to the theoretical reflectance data, and the resulting modeled reflectance was similar in magnitude to the measured values. Additionally, some scattered and diffracted light introduced from the optical and mechanical components in the instrument likely contributed to a background level of light present in the LTR images. In the end, we selected the minimum measured reflectance scaling method because it produced good fits to the data without making assumptions about the magnitudes of each contributing factor.

The measured thickness was returned by identifying the parameters that minimize the fit figure of merit (FOM, Equation 1).

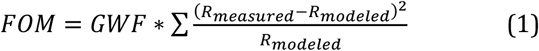

The fit FOM is based on Pearson’s Chi-square statistic, a normalized Method of Least Squares that ensures that larger reflectivity values do not dominate the fit, which is important for locating minimum reflectivity. To improve the fit, a 2D Gaussian weighting function centered on the reflectance null is applied to increase the influence of values closer to the minimum (Equation 2), where L and Q represent the widths of the major and minor axes of the elliptical gaussian, and *θ* determines the rotational orientation of the weighting function, *x*_0_ *is x*_*min*_, and*y*_0_ is *y*_*min*_.

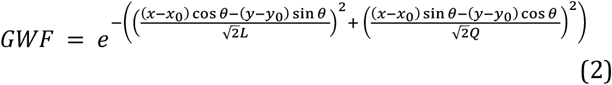

This method produced good agreement between the theoretical and measured data over the full measurement range (Figure 3c). This fitting procedure leveraged the full slope of the steep null reflectivity condition and was more robust than a simple peak finding algorithm. Importantly, data points that sample the reflectance properties around the null contain valuable information that can be used to uniquely identify thickness even when the depth of the observed null is limited by scattered light or substrate roughness. Shifts of 1 Å, 0.1 Å, and 0.01 Å in modeled thickness increase the fit FOM by 634%, 6%, and 0.02% respectively (Figure S3).

The performance of the LTR instrument was evaluated by measuring 35 individual substrates with SiO_2_ film thicknesses ranging from 1390 to 1465 Å using both LTR and spectroscopic ellipsometry. The average difference between LTR and ellipsometry SiO_2_ thickness measurements was 0.03%, and a linear regression returned a slope of 0.997 and y-intercept of 3.72 (Figure 4). This demonstrates impressive agreement between two independent thickness measurement techniques, both of which have their own degree of measurement uncertainty. The uncertainty reported by the J.A. Woollam software with each ellipsometry measurement was ∼0.2 Å. The measured LTR thickness deviation of 40 repeat measurements of a single location on a single substrate was 0.01 Å. Overall, LTR was accurate to within 1% of ellipsometry measurements, with a 3-sigma precision of 0.03 Å.

**Figure 4.**
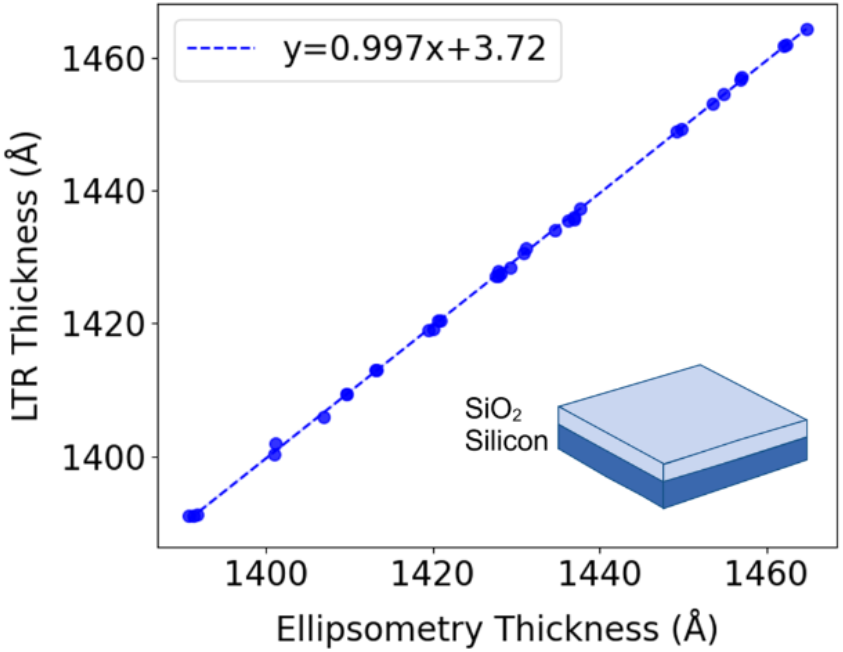
Comparison of measurements of 35 substrates with both LTR and ellipsometry.

We examined the ability of LTR to measure the thickness of protein films formed on the entire surface of a Si/SiO_2_ substrate. LTR measurements were made of separate substrates at different steps during the silane functionalization (glycidoxypropyl trimethoxysilane (GPTMS), adhesion layer, Figure 5a), probe deposition, and analyte binding process. There was a 40 Å thickness increase due to the covalent attachment of the IgG molecule to the GPTMS amine-reactive adhesion layer (Figure 5b). The binding of antibody to the IgG coated substrate caused an additional 35 Å increase in thickness (Figure 5c). These measurements of a monolayer of protein across the entire substrate surface demonstrate the movement of the null reflectivity condition in lambda theta space as a function of optical thickness and confirm the ability to observe changes in protein films due to antibody-antigen binding. We used high concentrations of IgG and anti-IgG (0.5 and 3 µg/mL respectively) for these experiments. Therefore, these protein films can be thought of as a closely packed monolayer of proteins on the surface of the substrate. The ∼75 Å total thickness change due to anti-IgG and IgG proteins is less than 6% of the thickness of the underlying SiO_2_ film (∼1400 Å). Therefore, we chose to approximate the protein film layer as an extension of the SiO_2_ layer in the characteristic matrix model. This is appropriate for biosensing applications where physical thickness is less important than relative signal change due to binding of analyte to probe.

**Figure 5.**
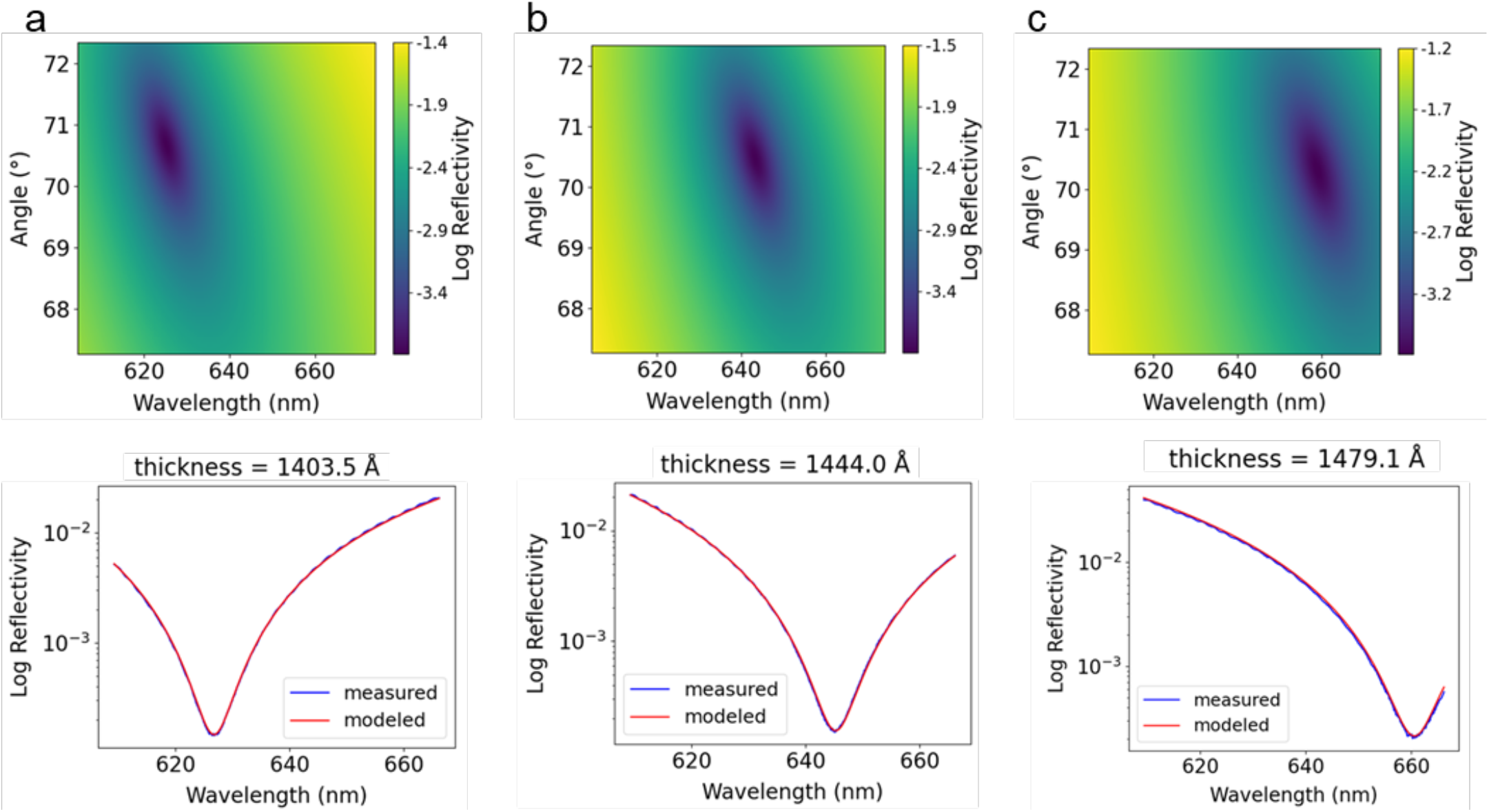
Biosensing capability of LTR with whole chip chemical and protein films. (a) LTR measurement of SiO_2_/Si substrate coated with amine-reactive silane. (b) LTR measurement of SiO_2_/Si substrate coated with amine-reactive silane and incubated in 0.5 µg/mL human IgG protein. (c) LTR measurement of SiO_2_/Si substrate coated with amine-reactive silane, incubated in 0.5 µg/mL IgG protein, and incubated in 3 µg/mL anti-IgG.

We next confirmed that LTR thickness measurements can be used to measure analyte concentration. A standard curve was constructed by arraying several substrates with anti-IL-6 analyte probes and anti-fluorescein isothiocyanate (α-FITC) control probes and incubating each substrate in different serially diluted concentrations of the analyte interleukin-6 (IL-6) overnight to allow the system to reach equilibrium. IL-6 is a 26 kDa protein that modulates cell-signaling pathways in the human immune system.^19^ It is normally present in human serum at very low concentrations less than 10 pg/mL. An elevated concentration (10s to 100s of pg/mL) of IL-6 in serum is a common biomarker of inflammation and a heightened immune response.^20^ Fluorescein iso-thiocyanate (FITC) is a molecule that is normally not present in human samples, so we use an antibody against FITC as a control probe to account for inter– and intra-substrate differences in the thickness of the underlying oxide layer.

The control substrate was exposed only to the assay diluent and was used to establish baseline thickness. It is the blank measurement. The analyte substrates were exposed to different concentrations of IL-6 mixed with diluent. The lower limit of detection (LLOD) is defined as the concentration of analyte at which the sensor measures a response that is significantly different than the blank. The limit of the blank (LOB) was calculated from the control values (Equation 3), where *μ* is the mean, and *σ* is the standard deviation. The LLOD was calculated from a combination of all low-concentration thickness values and standard deviations (Equation 4).^21, 22^

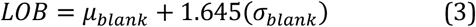

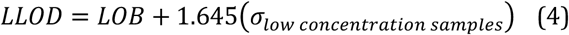

We calculated the LLOD of label-free IL-6 detection using a 4-parameter logistic regression (4PL) fit to relate anti-FITC-subtracted LTR thickness measurements to IL-6 concentration (Equation 5, Figure S4), where a is the minimum response, b is the Hill slope of the curve, c is the point of inflection, and d is the maximum response. The 4PL regression fits ligand binding response better than a 1:1 Langmuir binding isotherm in assays that deviate from perfectly independent binding events of analytes that are monovalent and homogeneous^23^ and generally outperforms other models to fit the full range of response data comprising the sigmoidal shape of biological binding assays.^24^

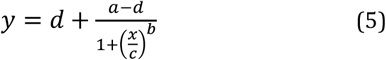

The calculated LLOD of 539 pg/mL (0.02 nM) is at the upper limit of the clinically relevant range for IL-6 in human serum, indicating that this assay in its current form would not be useful for measurements of IL-6 in blood samples. However, we note that this reflects a limitation of the binding affinity of the IL-6 capture antibody to its IL-6 target rather than a limitation of the instrument itself. To explore the effect of binding affinity on the sensitivity of this assay, we plotted LTR thickness change versus IL-6 analyte concentration and applied the 1:1 one-site Langmuir binding isotherm model.^25,17^ Thickness change measurements were calculated by subtracting anti-FITC-corrected thickness values on the control substrate (incubated only in diluent) from the anti-FITC-corrected thickness values on the analyte substrate (Equation 6). The Langmuir binding isotherm is an idealized model of 1:1 binding that describes the fractional coverage of binding sites as a function of analyte concentration (Equation 7).

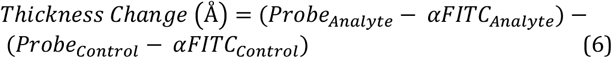

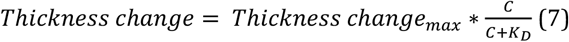

The system must be at dynamic equilibrium for this model to apply, and this assay was performed overnight at 4 °C to allow it to reach equilibrium. The *Thickness change*_*max*_ is the experimental maximum LTR thickness measurement for this IL-6 antibody-antigen pair assuming that analyte is bound to 100% of available probe sites. The dissociation constant (K_D_) represents the concentration at which half of the binding sites are occupied, and *C* is the Molar concentration of IL-6 analyte. The inflection point of the 4PL fit also represents the K_D_, but we chose to use the one-site Langmuir isotherm model because it offers an intuitive assessment of the effect of concentration and affinity on sensor response and is grounded in the principles of adsorption kinetics of molecules at a planar surface.

Using the one-site Langmuir binding isotherm model we calculated the equilibrium dissociation constant (K_D_) for the IL-6 binding pair in this experiment to be 8.4E-11 M. We then assessed the impact that a 10-fold increase or decrease in K_D_ would have on the binding isotherm (Figure 6).

**Figure 6.**
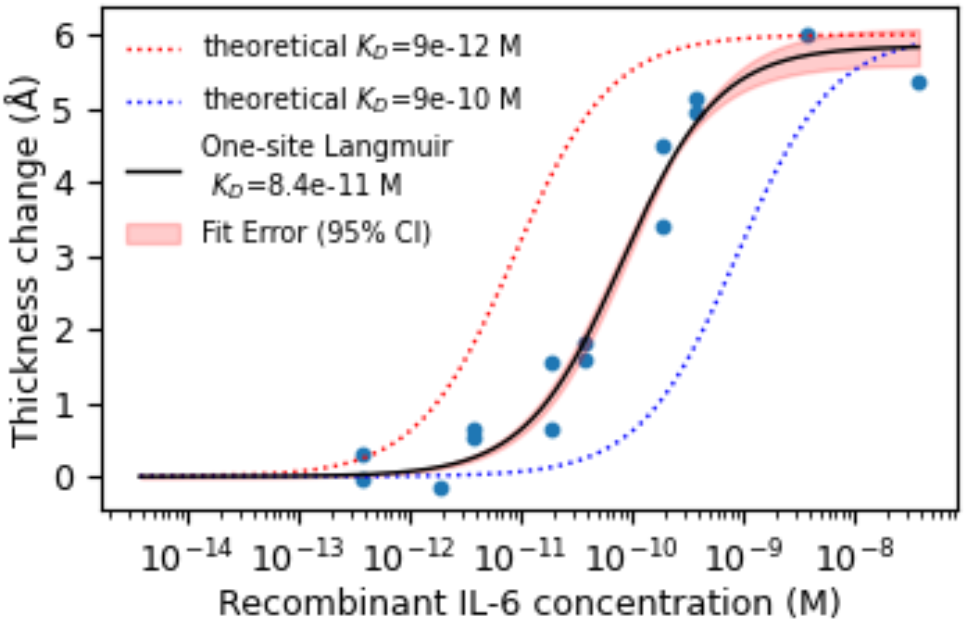
LTR measured thickness change relative to control substrate exposed only to diluent fit with a one-site Langmuir binding isotherm. An increase in binding affinity (decrease in K_D_) shifts the isotherm to improve detection at lower concentrations.

A lower K_D_ represents a stronger affinity between analyte and probe, which means more analyte remains bound to probe in equilibrium binding conditions at lower concentrations, presumably increasing the measurable signal at low concentrations. We further considered the effect of the binding pair K_D_ on assay sensitivity by calculating the expected LTR thickness change at several K_D_ values for 1 pg/mL of analyte (Table 1). The inability to detect a 0.0025 Å thickness change at 1 pg/mL for this IL-6 binding pair (K_D_ = 8.4E-11) is consistent with the 3*σ* resolution limit of the instrument (0.03 Å). If we could employ a higher affinity probe (K_D_ = ≤ 6E-12 M) in this assay, then the thickness change due to 1 pg/mL of IL-6 could be measured with LTR assuming no other sources of variation. However, differences between biological replicates would likely continue to limit the LLOD because this non-instrumental source of error has a significant contribution to the LLOD calculation.

**Table 1.** Theoretical thickness change required to measure 1 pg/mL of analyte at different binding pair (probe and analyte) K_D_ values. If the LTR thickness change resolution of 0.03 Å was the only factor governing the LLOD, then the assay could detect the thickness change due to 1 pg/mL of IL-6 binding to probe when the binding pair K_D_ is ≤ 6E-12 M.

| $K_D$ | 9E-14 | 9E-13 | 4E-12 | 5E-12 | 6E-12 | 7E-12 | 8E-12 | 9E-12 | 9E-11 | 9E-10 |
| --- | --- | --- | --- | --- | --- | --- | --- | --- | --- | --- |
| Calculated thickness change (Å) at 1 pg/mL of IL-6 | 1.720 | 0.237 | 0.076 | 0.057 | 0.046 | 0.033 | 0.029 | 0.0246 | 0.0025 | 0.0002 |

The motivation for designing and constructing this LTR instrument was a desire to overcome the limitations of AIR when initial probe thickness is not optimized to the antireflective baseline condition. To demonstrate the improvement that LTR offers over AIR we compared measurements of an array of probes rejected from a previous AIR study because it was fabricated on a “too thin” substrate. This array consists of 21 proteins produced by *Staphylococcus aureus* (6 replicate spots of each) and 71 replicate anti-FITC correction probes around the border (Figure S5). The *S. aureus* proteins are found on the surface of the bacteria or are secreted into the environment and are known to have immunomodulatory effects^26^ including production of anti-*S. aureus* antibodies with lock-and-key specificity. A version of this multiplex array (built on substrates with optimal baseline thickness for AIR) was used in previous work to quantify the human antibody response to *S. aureus* bacteria in the contexts of infection or related disease.^27,28^ In brief, the array consisted of proteins representing 6 broadly defined functional classes:

1. Iron acquisition proteins (iron-regulated surface determinant proteins IsdA, IsdB, IsdH)
2. Cell division and cell wall proteins (autolysin domains glucosaminidase [Gmd] and amidase [Amd], immunodominant staphylococcal antigen A [IsaA])
3. Biofilm formation proteins (clumping factor A [ClfA] and bone sialoprotein-binding proteins [BsBp])
4. Immune evasion proteins (chemotaxis inhibitory protein of *Staphylococcus* [CHIPS]; staphylococcal complement inhibitory protein [SCIN])
5. Superantigens (staphylococcal enterotoxin A [SEA], B [SEB], and C [SEC]; staphylococcal enterotoxin-like G [SE*l*G], I [SE*l*I], Q [SE*l*Q], X [SE*l*X], and toxic shock syndrome toxin-1 [TSST-1])
6. Cytotoxins (leukocidin F [LukF] and S [LukS], and alpha-hemolysin [Hla]).

We were able to identify the probes that are too thin to be accurately measured by AIR by measuring the LTR thickness of each probe on the control substrate (Figure 7). This is the first time that we have been able to accurately measure the baseline probe thickness on these “too-thin” arrays.

**Figure 7.**
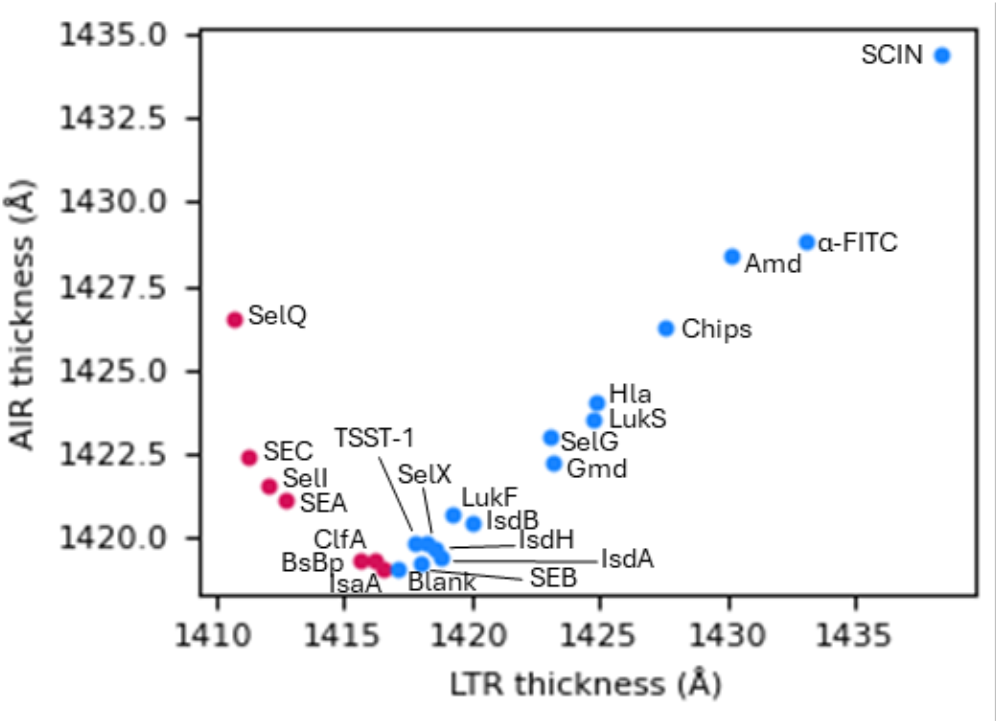
LTR provides accurate thickness measurements for all *S. aureus* probes on a sub-optimal AIR array. The probes surrounded by the red boxes are the probes that are too thin for accurate AIR measurements (due to using the positive quadratic root to return thickness by default) but were easily measured with LTR.

To further demonstrate the ability for LTR to make accurate thickness change measurements regardless of initial probe thickness, we incubated the array in a 1:250 diluted serum from an individual with a culture-confirmed *S. aureus* infection. The measured thickness change of the “thin” probes as measured by both AIR and LTR is shown in Figure 8. LTR was able to rescue the functionality of these probes and produce robust measurements of thickness change due to bound antibody. The most dramatic improvement in binding response was observed for the SEA, SEC, SelI, and SEQ antigens.

**Figure 8.**
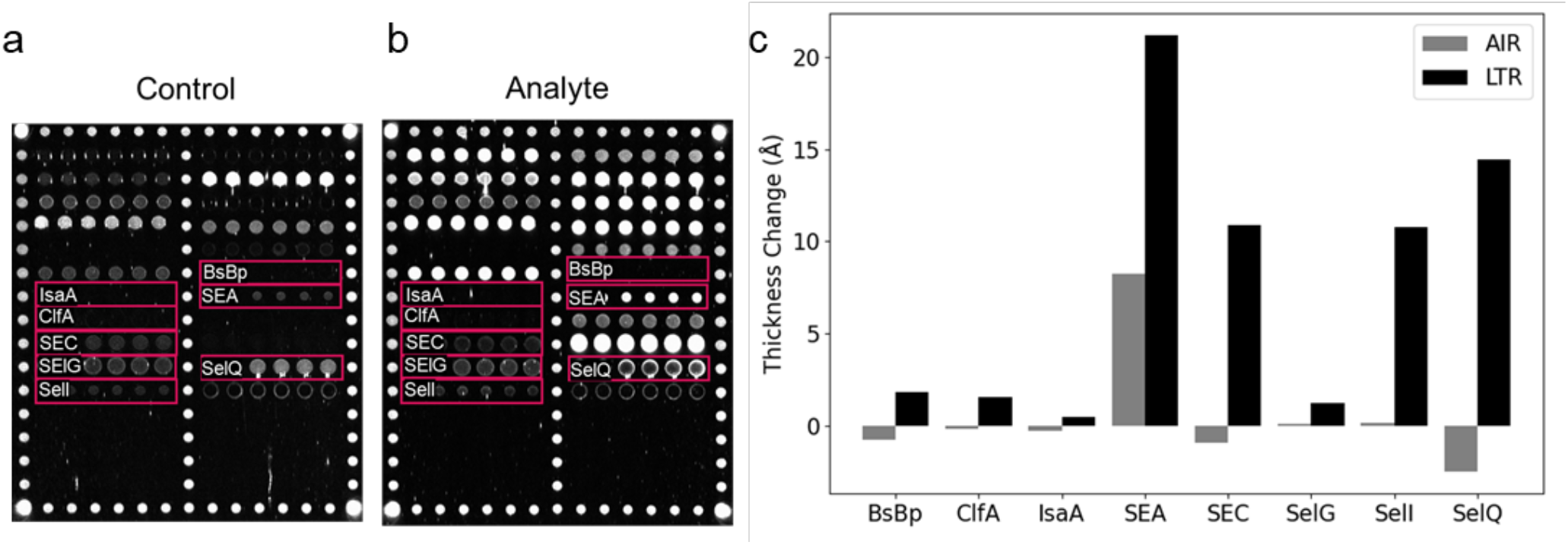
LTR returns accurate binding measurements for thin probes and “rescues” probes that AIR cannot measure accurately. (a) AIR image of Baseline probe thickness. The control chip is only exposed to assay diluent. (b) AIR image of a substrate exposed to serum from an individual with a culture-confirmed *S. aureus* infection. (c) AIR and LTR thickness change measurements for probes that are on the left of the AIR reflectivity vs thickness parabola and labeled in AIR images. LTR thickness change measurements are not limited by baseline thickness

## Conclusions

The key technical advance of LTR is the simultaneous illumination of a thin film substrate over a range of wavelengths and AOIs with the ability to resolve angular and spectral content of the reflected light along two orthogonal dimensions concurrently. This enables LTR to identify the null reflectivity condition over a range of film thicknesses without scanning through angle or wavelength.

LTR fundamentally measures optical thickness. For an absolute physical thickness measurement, an external measurement of refractive index would be required. This is not necessary when using LTR as a biosensor, since the metric of interest is thickness change due to analyte binding to probe molecules, and a relative thickness measurement is appropriate. Furthermore, the protein refractive index varies according to the illumination wavelength as well as amino acid content, concentration, temperature, and extent of hydration^29^, which would be difficult to measure and model for each analyte and probe on the array.

While the ability of LTR to measure protein binding on planar arrays is encouraging, we have considered avenues for further improvement. The sensitivity at low concentrations could be enhanced by selecting probes with higher affinity (lower K_D_) for the analyte of interest. Another approach would be to include a sandwich antibody or other mass-based amplification strategies to increase the thickness at low analyte concentrations. The measurement sensitivity also depends on the variation between biological replicates because the standard deviations of the blank and the low concentration samples are fundamental to the LLOD calculation. There are differences in base oxide thickness both between control and analyte chips as well as across the surface of a single chip. Hyper-localized corrections for baseline thickness variations could decrease the variability between replicates. This would either mean using an adjacent α-FITC probe or sampling the background area immediately next to each analyte probe for inter– and intra-chip correction rather than using only a single α-FITC measurement or averaging values across a substrate.

The thickness resolution of the LTR instrument itself may be enhanced by deepening the null reflectivity condition. We assessed several sources of noise in the system including scattered light, temperature effects, and other factors affecting finite resolution limits. The variation in the repeated measurements appears to be a function of temperature changes and lamp output fluctuations over the course of the measurement window. Localized heating at the illumination region as well as increased ambient temperature in the instrument box from heat dispelled from the sensor electronics could cause changes in the optical thickness of the film over time. Physical layer thickness and material index of refraction are both temperature dependent which can lead to drifts in the measured optical thickness.

It is important to consider, however, that ultra-low LLODs are not the only metric for evaluating biosensor performance. Other considerations are ease of use, accuracy, multiplexing capability, and dynamic range, which are all strengths of LTR. The Si/SiO_2_ substrate is large enough to fit hundreds of probes, and the 100 Å measurement range eliminates the need for multiple sample dilutions. Since LTR is a label-free measurement, mass-based amplification techniques could target low concentration analytes without negatively affecting high-concentration signals. While LTR requires scanning of the substrate to image an entire array (unlike AIR, which acquires a single image of the entire substrate), programmable translational stages could allow the task to be accomplished simply and rapidly. LTR returns a single thickness measurement and does not require precise tuning of baseline probe thickness, which enables highly accurate measurements with relaxed array manufacturing requirements. This will be particularly helpful in our ongoing work studying the antibody response to S. aureus infection. Other potential applications include the study, diagnosis, and treatment of autoimmune^30^, neuroinflammatory^31^, and infectious disease^32^. Biomarker panels are typically more effective at diagnosing disease than single-plex assays,^33^ and they provide more information from a single sample in scientific studies. LTR could be used to measure concise panels consisting of biomarker antigens, cytokines, and antibodies. This type of “mixed assay” is possible because LTR is a label-free measurement, and the sensing range spans several logs of concentration while requiring only a single sample dilution. Furthermore, the sensing range of LTR is controlled by the range of wavelengths and angles present in the illumination, which can be expanded if needed.

We developed LTR to address the challenging workflow of current reflectometric methods, in particular the stringent requirements for baseline probe thickness to achieve the antireflective condition when using AIR. The prototype LTR instrument measured SiO_2_ film thickness accurate to within 1% of ellipsometry and demonstrated impressive 0.01Å precision. It measures SiO_2_ and protein film thickness over a 100 Å range, representing concentrations ranging from < 1 ng/mL of small IL-6 protein to 3 µg/mL of larger IgG and anti-IgG proteins. LTR was able to recover antibody binding information from a *S. aureus* antigen array that was not optimized for AIR. Specific antibodies against *S. aureus* proteins are present at concentrations greater than 1 µg/mL in human serum^34^. This large sensing range allows multiple protein types to be measured simultaneously from a single sample dilution. The ability to make accurate measurements of multiplexed biological arrays while loosening the probe deposition requirements will enable us to use this technique to study a wider set of biomarkers.

## Materials and Methods

### LTR Instrument Calibration

The LTR optics were arranged to sort and direct photons to specific locations at the detector plane based on their wavelength and angle of incidence at the biosensor surface. Wavelength calibration was established using a Neon spectral calibration lamp that produces 15 detectable emission lines spanning 607 to 671nm. Each of these lines provided a unique wavelength position and spectral resolution measurement at a different location across the image. The spectral resolution varies over the field of view and ranges from 0.6 nm to 1.0 nm full width at half maximum. The optical design optimized spectral resolution by balancing the angular width of the slit aperture compared to the spread of angles introduced by diffraction as light passed through the slit. Wavelength values at pixel positions in between calibration points were inferred using a cubic spline interpolation.

A value for the angular span across a single pixel (Δ*θ*/Δpixel) was established by locating the edges of the illumination field and relating the cutoff positions on the image to the angular extent of the system aperture stop. A pinhole placed in the center of the optical axis generates a set up image that identifies the 70° reflectance AOI based on the geometry of the nominal opto-mechanical design. Together, this information was used to generate a linear relationship between AOI and pixel position. In practice these angular calibration techniques were used as a starting point for analysis and were later refined empirically by varying the values (+/-5%) to improve the goodness of fit of the LTR images.

Absolute reflectance measurements for each pixel in the LTR image were made by first recording a flat field calibration that baselines the relative signal levels using a substrate with a known reflectance. A Si/SiO_2_ substrate coated with 300 nm of Aluminum and 3 nm of SiO2 by E-beam physical vapor deposition served as a reflectance standard by with is nominal reflectivity, R_FF_(λ,*θ*), calculated from first principles. The signal present in a given pixel, ADU_i_(x,y) was determined by Equation 8 where P is the lamp emission given in Photons/second, R is the reflectivity of the substrate, T is the system transmission, including diffraction and optic coatings, QE is the quantum efficiency of the detector (CCD e-/ photon), G is camera gain given in (ADU / CCDe-), and t is the image exposure duration.

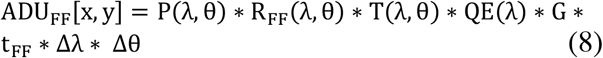

A similar expression determined the signals observed when a functionalized Si/SiO_2_ chip was used. The absolute reflectance of the LTR data was calculated using a ratio of these two images. Assuming the instrument sensitivity parameters and lamp output power remain constant over time, many parameters cancel, and the reflectance is given by Equation 9.

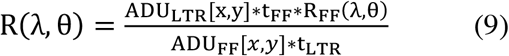

An external shutter controls the lamp exposure duration and is adjusted such that the sensor well capacity is nearly full in the brightest regions of the image. Typical exposure durations are 200 ms for flat field calibrations and 10 seconds for LTR images.

### Substrate Preparation

Polished 300 mm silicon wafers with 140 to 150 nm of thermally grown silicon dioxide were fabricated by SUNY Polytechnic Institute, diced into 5 × 5.7 mm substrates, and binned by oxide thickness according to measurements made by a J.A. Woolam M2000 ellipsometer. Substrates were taken from a range of binned thicknesses, cleaned in piranha solution (3 parts sulfuric acid to 1 part 30% hydrogen peroxide) for 30 minutes, washed 3x in Nanopure water, and dried with a stream of N_2_ gas prior to LTR measurements. The substrates used for protein measurements were selected to have similar oxide thickness and washed to remove dicing debris (7:3 solution of ethanol and 10 M NaOH) for 30 minutes, then etched in hydrofluoric acid to reach ∼1400 Å, and rinsed with water and dried with N_2_ gas before being chemically functionalized with ∼5 Å of amine-reactive (3-glycidoxypropyl) trimethoxysilane (GPTMS(Sigma Aldrich 440167)) in a plasma process chemical vapor deposition system (YES 1224P) at the University of Rochester Integrated Nanosystems Center.

### Protein films across entire substrate surface

Surface protein films were grown by incubating entire substrates overnight at room temperature in a 0.5 µg/mL solution of human IgG whole molecule protein (Rockland 009-0102) and phosphate buffered saline (PBS: 137 mM NaCl, 2.7 mM KCl, 10 mM Na_2_HPO_4_, 1.76 mM KH_2_PO_4_·H_2_O, pH 7.4) at 300 RPM on an orbital shaker. Substrates were next placed in assay wash buffer (AWB: mPBS with 0.005% Tween-20, pH 7.2) for 5 minutes before incubating in (3 µg/mL) anti-IgG protein (Rockland 609-701-123) solution in PBS for 1 hour at room temperature. Substrates were removed from each step of this process for LTR measurements. All substrates were washed with Nanopure water and dried with a stream of N_2_ gas before being measured.

### AFM measurements

A NTMDT AFM Microscope was used at the University of Rochester Integrated Nanosystems center in semi-contact tapping mode at a 10 µm/second scan rate over a 5 µm^2^ area. Image post-processing and power spectral density analysis was accomplished using open source Gwyddion software^35^.

### Arraying protein probes onto substrates

Substrates with arrayed probes first went through the above-described base wash, HF etch, and GPTMS functionalization process before having proteins solutions deposited by an SX sciFLEXARRAYER (Scienion A.G.; Berlin, Germany) with a PDC70 capillary with type 4 coating. The protein probe mixtures comprising the microarray were pipetted into individual wells of a 384-well plate. Anti-IL-6 (Biolegend 501125) was diluted in PBS to 400 µg/mL. Anti-FITC (Rockland 600-101-096) was dialyzed to remove sodium azide by using 20 kDa molecular weight cutoff Slide-a-Lyzer dialysis cups floating in a beaker of PBS stirred slowly with a magnetic stir bar) for 1.5 h at RT before dilution in PBS to a concentration of 500 µg/mL for arraying. Recombinant *S. aureus* antigens were selected and produced with a biotin tag as described previously^36^. The *S. aureus* antigen formulations are detailed in Figure S6. All probe solutions contained 2% trehalose as an additive to improve spot morphology. Substrates were mounted onto adhesive strips and placed inside the SX arrayer, which was set to 85% ± 4% relative humidity. Droplets were between 300 and 350 pL in volume, as measured by the instrument. After completion of the arraying process, the chips remained in the humidified chamber overnight to ensure covalent attachment of the protein probes to the GPTMS-functionalized substrates.

### Blocking the background of arrayed substrates

All incubations were performed with stabilized microarrayed chips mounted on adhesive combs inserted into individual wells of a 96-well plate on an orbital shaker at 300 rpm. The probe-arrayed chips were removed from the humidified chamber and immediately placed into wells of a 96-well plate, each containing 300 *μ*L of 50 mM NaOAc, pH 5.0, for 5 min to prevent smearing of any unadsorbed probes onto nearby areas of the amine-reactive chip surface. The chips were then incubated in 1% bovine serum albumin (BSA: Rockland BSA-50) in NaOAc, pH 5.0, for 30 min to block the background from nonspecific adsorption and then transferred to wells containing a second blocking solution of 20% fetal bovine serum (FBS: Gibco A5670701) in assay wash buffer (AWB: mPBS with 0.005% Tween-20, pH 7.2) for 30 min. All incubation steps were performed at RT with orbital shaking (300 RPM).

### Analyte incubations

The assay diluent in all cases was Enhanced Assay Buffer (EAB: 20 mM Tris Base, 250 mM NaCl, 250 mM KCl, 3% w/v propylene glycol, 0.125% Triton X-100, and 1% w/v BSA) with 20% FBS. Each substrate for the IL-6 standard curve was incubated in serially diluted (10000, 5000, 1000, 500, 100, 50, and 10 pg/mL) recombinant human IL-6 (Biolegend 570804) overnight at 4 °C shaking at 420 RPM. Substrates were then washed in AWB and Nanopure and dried with a stream of N_2_ gas. Substrates for *S. aureus* antibody detection were incubated in a 1:250 dilution of human serum from an individual with a culture-confirmed *S. aureus* infection overnight at 4 °C shaking at 420 RPM, washed in AWB for 5–10 min, then exposed to a 1 h RT incubation with an Fcγ-specific anti-IgG (AffiniPure 109-005-008) to amplify the response. This anti-IgG amplification step was performed to maintain consistency with previous AIR assays performed using similar *S. aureus* arrays. Chips underwent a final incubation in AWB for 10 min before being rinsed in Nanopure water and dried under a stream of nitrogen gas. Control chips underwent the same process but were only exposed to the assay diluent during the primary incubation step.

### AIR image analysis

AIR images were analyzed in ImageJ. Regions of interest were drawn around individual probe spots to measure the median pixel intensity. These intensity values were converted to thickness using the equation of the parabolic fit of the AIR vs. ellipsometry calibration. The piranha-washed Si/SiO2substrates with a range of oxide thickness were also measured with AIR to construct the AIR vs ellipsometry calibration curve. The AIR intensity value depends on the CCD exposure time, so the exposure time of the image was matched to the calibration curve exposure time. Care was taken to avoid measuring probe spots that were over-exposed, so multiple AIR images were collected over a range of exposure times. Measurements were taken from images where median intensity was 30,000 to 40,000 counts.

## ASSOCIATED CONTENT

### Supporting Information

Limitations of AIR, reflectivity vs thickness curve; AFM micrograph of a Si/SiO_2_ substrate with power spectral density analysis; Fit FOM vs modeled thickness; 4-parameter logistic fit of IL-6 LTR measurements; *S. aureus* probe locations on array; *S. aureus* antigen formulations. (PDF) “This material is available free of charge via the Internet at http://pubs.acs.org.”

## AUTHOR INFORMATION

## Author Contributions

The manuscript was written through contributions of all authors. All authors have given approval to the final version of the manuscript. **Alanna Klose:** Conceptualization, Methodology, Software, Investigation, Writing-Original draft, review, and editing, Funding acquisition **Joseph Katz:** Conceptualization, Methodology, Software, Investigation, Validation, Writing-Original draft, Review and Editing. **Robert Boni:** Conceptualization, Software, Methodology, Writing-Review and Editing. **David Nelson:** Conceptualization, Mechanical design. **Benjamin Miller:** Conceptualization, Supervision, Writing – Review and Editing, Funding acquisition.

## ACKNOWLEDGMENT

The authors thank Dr. Patrick Schievert for generously sharing *S. aureus* enterotoxins, and Dr. Gowri Muthukrishnan and Dr. Cheryl Ackert Bicknell for providing a human serum sample from an individual with an *S. aureus* spine infection. This sample was collected at the University of Colorado under an approved Institutional Review Board protocol #20-2946. They also thank Dr. Dustin Froula at the University of Rochester Laboratory for Laser Energetics for supporting this project and Timothy Ilardo for assisting with instrument assembly.

## ABBREVIATIONS

LTR: Lambda Theta Reflectometry
OI-RD: Oblique-Incidence Reflectivity Difference
RIfS: Reflectometric Interference Spectroscopy
BLI: Biolayer Interferometry
SRIB/IRIS: Spectral Reflectance Imaging Biosensor/Interferometric Reflectance Imaging Sensor
AIR: Arrayed Imaging Reflectometry
AOI: angle of incidence
AFM: atomic force microscopy
FOM: figure of merit
GPTMS: glycidoxypropyl trimethoxysilane
FITC: fluorescein isothiocyanate
IL-6: interleukin-6
LLOD: lower limit of detection
LOB: limit of the blank
4PL: four parameter logistic
K_D_: dissociation constant
IsdA,B,H: Iron-regulated surface determinant protein A,B,H
Gmd: glucosaminidase
Amd: aminidase
IsaA: immunodominant staphylococcal antigen A
ClfA: clumping factor A
BsBp: bone sialoprotein-binding protein
CHIPS: chemotaxis inhibitory protein of *Staphylococcus*
SCIN: staphylococcal complement inhibitory protein
SEA,B,C: staphylococcal enterotoxin A,B,C
Se*l*G,I,Q,X: staphylococcal enterotoxin-like G,I,Q,X
TSST-1: toxic shock syndrome toxin-1
LukF: leukocidin F
LukS: leukocidin S
Hla: alpha-hemolysin
PBS: phosphate buffered saline
AWB: assay wash buffer
BSA: bovine serum albumin
EAB: enhanced assay buffer

## TOC image

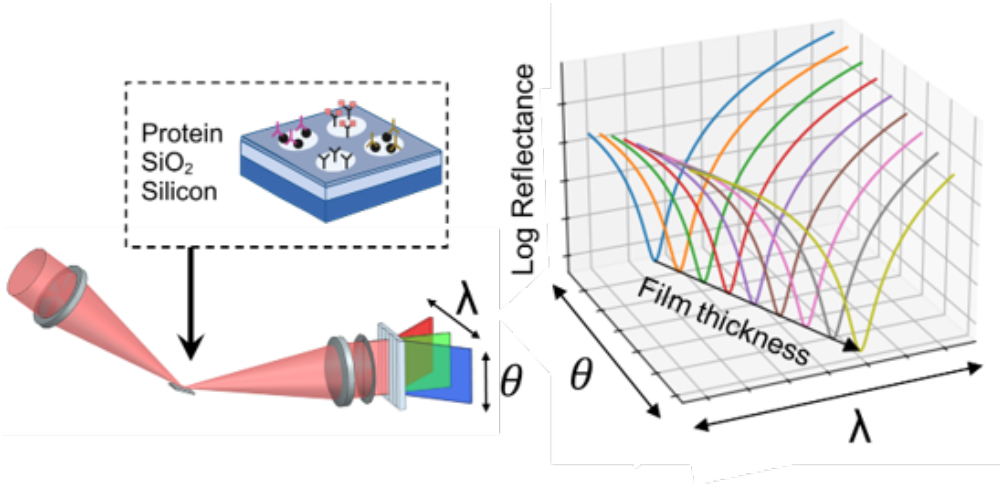

