## Supplemental figures for "Lambda Theta Reflectometry: a new technique to measure optical film thickness applied to planar protein arrays"

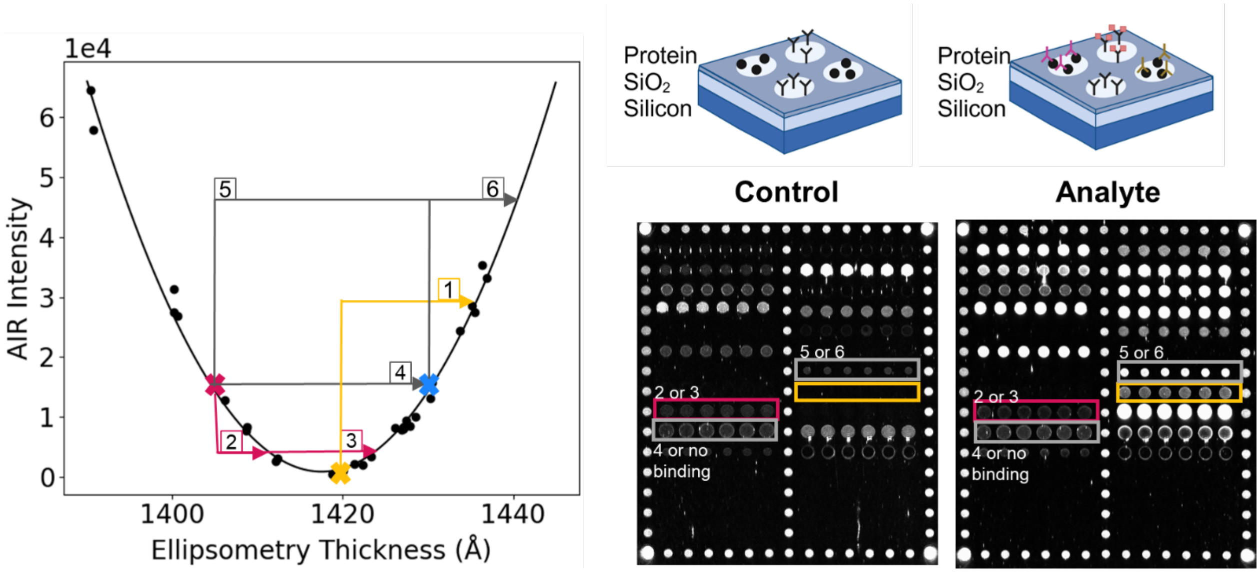


**Figure S1**. The AIR reflectivity vs thickness curve is parabolic. When converting reflectivity to thickness the positive root is taken by default assuming that probes are engineered to start at the antireflective condition and increase in thickness as analyte binds. When baseline probe spot thickness is thinner or thicker than the optimal condition, then it is unclear whether they are starting on the left or the right of the AIR reflectivity vs thickness curve. Optimal probe deposition thickness for AIR is shown by yellow x (scenario #1). Problematic probe deposition scenarios are indicated by the magenta and cyan x's. A thin probe that decreases in intensity after binding analyte could represent two different thickness changes (scenario #2 or #3). An unoptimized probe that doesn't change in intensity could represent either 0 Å or ~20 Å of thickness change (scenario #4). Another unoptimized probe that increases in reflectivity could indicate scenario #5 or #6.

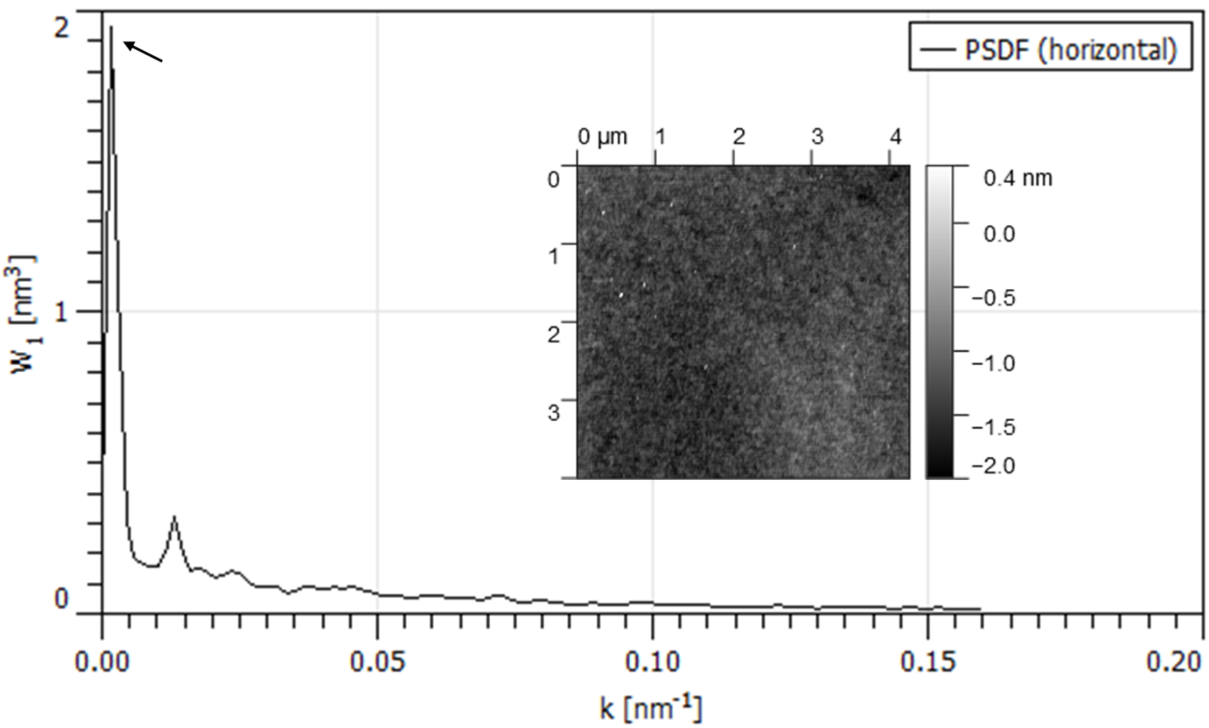


**Figure S2.** AFM micrograph of an ~4 µm$^2$ area (10 µm/second scan rate) of a $Si/SiO_2$ substrate. The horizontal power spectral density function shows a high frequency of height variation on a length scale of 0.0015 nm-1, or 684 nm (indicated by arrow). This frequency is larger than the wavelengths of light used by LTR and these height differences could result in the superposition of null reflectivity values.

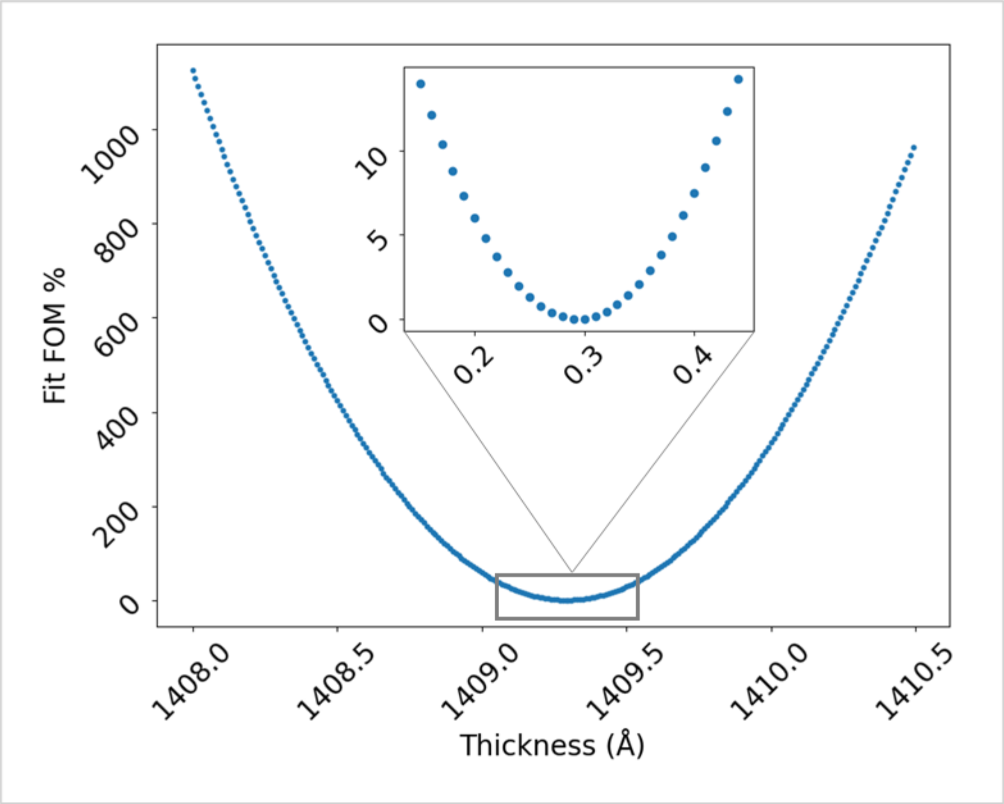


**Figure S3.** Fit FOM vs modeled thickness. Shifts of 1 Å, 0.1 Å, and 0.01 Å in modeled thickness increase the fit FOM by 634%, 6%, and 0.02% respectively.

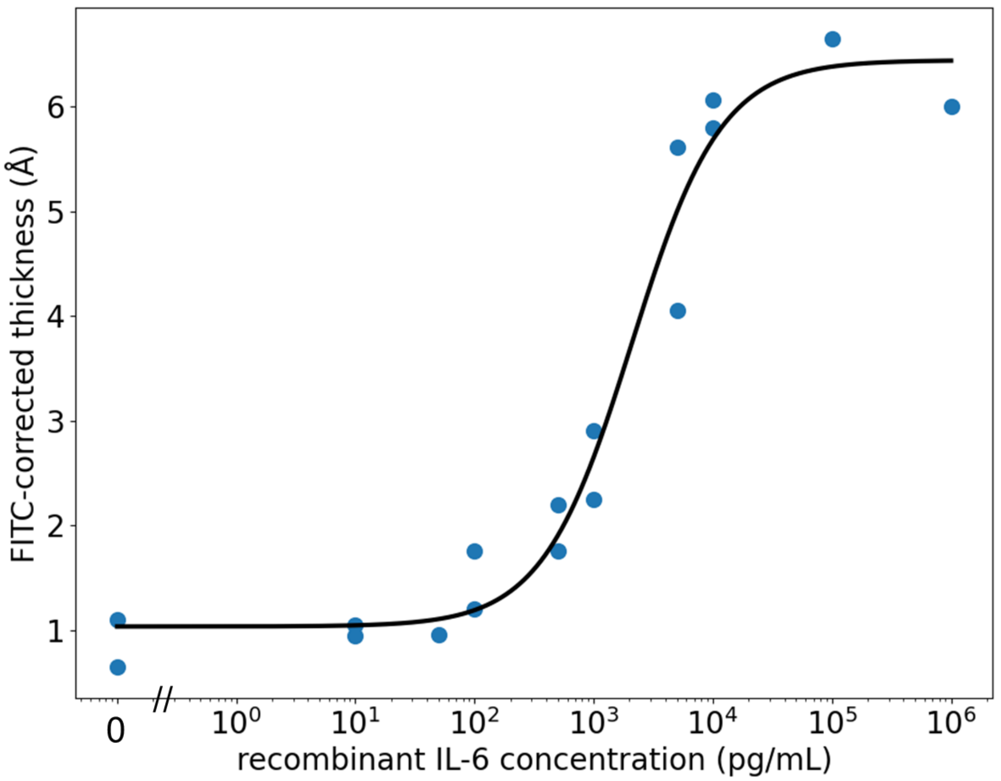


**Figure S4.** 4-parameter logistic fit of FITC-corrected LTR thickness measurements of IL-6. This fit was used to return the LLOD of 539 pg/mL for this assay.

| | | | | | | | | | | | | | | |
|---|---|---|---|---|---|---|---|---|---|---|---|---|---|---|
| hIgG | α-FITC | α-FITC | α-FITC | α-FITC | α-FITC | α-FITC | α-FITC | α-FITC | α-FITC | α-FITC | α-FITC | α-FITC | α-FITC | hIgG |
| α-FITC | IsdB | IsdB | IsdB | IsdB | IsdB | IsdB | α-FITC | IsdH | IsdH | IsdH | IsdH | IsdH | IsdH | α-FITC |
| α-FITC | Gmd | Gmd | Gmd | Gmd | Gmd | Gmd | α-FITC | SCIN | SCIN | SCIN | SCIN | SCIN | SCIN | α-FITC |
| α-FITC | Hla | Hla | Hla | Hla | Hla | Hla | α-FITC | IsdA | IsdA | IsdA | IsdA | IsdA | IsdA | α-FITC |
| α-FITC | Amd | Amd | Amd | Amd | Amd | Amd | α-FITC | CHIPS | CHIPS | CHIPS | CHIPS | CHIPS | CHIPS | α-FITC |
| α-FITC | Blank | Blank | Blank | Blank | Blank | Blank | α-FITC | LukF | LukF | LukF | LukF | LukF | LukF | α-FITC |
| α-FITC | LukS | LukS | LukS | LukS | LukS | LukS | α-FITC | BsBp | BsBp | BsBp | BsBp | BsBp | BsBp | α-FITC |
| α-FITC | IsaA | IsaA | IsaA | IsaA | IsaA | IsaA | α-FITC | SEA | SEA | SEA | SEA | SEA | SEA | α-FITC |
| α-FITC | ClfA | ClfA | ClfA | ClfA | ClfA | ClfA | α-FITC | SEB | SEB | SEB | SEB | SEB | SEB | α-FITC |
| α-FITC | SEC | SEC | SEC | SEC | SEC | SEC | α-FITC | TSST1 | TSST1 | TSST1 | TSST1 | TSST1 | TSST1 | α-FITC |
| α-FITC | SelG | SelG | SelG | SelG | SelG | SelG | α-FITC | SelQ | SelQ | SelQ | SelQ | SelQ | SelQ | α-FITC |
| α-FITC | SelI | SelI | SelI | SelI | SelI | SelI | α-FITC | SelX | SelX | SelX | SelX | SelX | SelX | α-FITC |
| α-FITC | Blank | Blank | Blank | Blank | Blank | Blank | α-FITC | Blank | Blank | Blank | Blank | Blank | Blank | α-FITC |
| α-FITC | Blank | Blank | Blank | Blank | Blank | Blank | α-FITC | Blank | Blank | Blank | Blank | Blank | Blank | α-FITC |
| α-FITC | Blank | Blank | Blank | Blank | Blank | Blank | α-FITC | Blank | Blank | Blank | Blank | Blank | Blank | α-FITC |
| α-FITC | Blank | Blank | Blank | Blank | Blank | Blank | α-FITC | Blank | Blank | Blank | Blank | Blank | Blank | α-FITC |
| α-FITC | α-FITC | α-FITC | α-FITC | α-FITC | α-FITC | α-FITC | α-FITC | α-FITC | α-FITC | α-FITC | α-FITC | α-FITC | α-FITC | α-FITC |

**Figure S5.** Grid defining the *S. aureus* proteins comprising each spot on the *S. aureus* array.

| | # drops | Probe | Undialyzed Concentration (µg/mL) | 1:4 dilution | 1:8 dilution | printed conc. (µg/mL) | volume of probe (µg) | volume of 9 µM avidin (µL) | volume of 7.4 pH PBS (µL) | volume of 25% trehalose (µL) |
|---|---|---|---|---|---|---|---|---|---|---|
| 1 | 1 | anti-FITC biotin | 1077 | | | 200 | 0.9 | 0.5 | 3.2 | 0.4 |
| 2 | 1 | human IgG | 5000 | | | 800 | 0.8 | 0 | 3.8 | 0.4 |
| 3 | 2 | ClfA | 300 | | | 120 | 4.1 | 0.5 | 0.0 | 0.4 |
| 4 | 2 | SEA biotin | 951 | | | 400 | 2.1 | 0.5 | 2.0 | 0.4 |
| 5 | 2 | SEB biotin | 954 | | | 400 | 2.1 | 0.5 | 2.0 | 0.4 |
| 6 | 2 | SEC biotin | 951 | | | 400 | 2.1 | 0.5 | 2.0 | 0.4 |
| 7 | 2 | TSST-1 biotin | 1306 | | | 800 | 3.1 | 0.5 | 1.0 | 0.4 |
| 8 | 2 | SelG biotin | 1218 | | | 900 | 3.7 | 0.5 | 0.4 | 0.4 |
| 9 | 2 | SelQ biotin | 500 | | | 410 | 4.1 | 0.5 | 0.0 | 0.4 |
| 10 | 2 | Sel I biotin | 487 | | | 200 | 2.1 | 0.5 | 2.0 | 0.4 |
| 11 | 1 | Sel X biotin | 1814 | | | 500 | 1.4 | 0.5 | 2.7 | 0.4 |
| 12 | 1 | IsdB | 4700 | | 587.5 | 200 | 1.7 | 0.5 | 2.4 | 0.4 |
| 13 | 1 | IsdH | 1300 | | | 200 | 0.8 | 0.5 | 3.3 | 0.4 |
| 14 | 1 | Gmd | 1000 | | | 200 | 1.0 | 0.5 | 3.1 | 0.4 |
| 15 | 1 | SCIN | unknown | | | | 1 | 1 | 7.2 | 0.8 |
| 16 | 1 | Hla | 3800 | 950 | | 400 | 2.1 | 0.5 | 2.0 | 0.4 |
| 17 | 1 | IsdA | 3100 | 775 | | 400 | 2.6 | 0.5 | 1.5 | 0.4 |
| 18 | 1 | Amd | 9000 | | 1125 | 200 | 0.9 | 0.5 | 3.2 | 0.4 |
| 19 | 1 | Chips | 2300 | 575 | | 400 | 3.5 | 0.5 | 0.6 | 0.4 |
| 20 | 1 | LukS | 3500 | 875 | | 200 | 1.1 | 0.5 | 3.0 | 0.4 |
| 21 | 1 | BsBp/SdrE | 600 | | | 400 | 3.3 | 0.5 | 0.8 | 0.4 |
| 22 | 1 | IsaA | 480 | | | 390 | 4.1 | 0.5 | 0.0 | 0.4 |
| 23 | 1 | LukF | 1624 | | | 200 | 0.6 | 0.5 | 3.5 | 0.4 |

**Figure S6.** Antigen formulations for printing the *S. aureus* antigens into the array on the $Si/SiO_2$ substrate.
